# Cutting through the noise: reducing bias in motor adaptation analysis

**DOI:** 10.1101/2020.11.25.397992

**Authors:** Daniel H. Blustein, Ahmed W. Shehata, Erin S. Kuylenstierna, Kevin B. Englehart, Jonathon W. Sensinger

## Abstract

During goal-directed movements, the magnitude of error correction by a person on a subsequent movement provides important insight into a person’s motor learning dynamics. Observed differences in trial-by-trial adaptation rates might indicate different relative weighting placed on the various sources of information that inform a movement, e.g. sensory feedback, control predictions, or internal model expectations. Measuring this trial-by-trial adaptation rate is not straightforward, however, since externally observed data are masked by noise from several sources and influenced by inaccessible internal processes. Adaptation to perturbation has been used to measure error adaptation as the introduced external disturbance is sufficiently large to overshadow other noise sources. However, perturbation analysis is difficult to implement in real-world scenarios, requires a large number of movement trials to accommodate infrequent perturbations, and the paradigm itself might affect the movement dynamics being observed. Here we focus on error adaptation during unperturbed and naturalistic movements. With increasing motor noise, the conventional estimation of trial-by-trial adaptation increases, a counterintuitive finding that is the consequence of systematic bias in the estimate due to noise masking the learner’s intention. We present an analytic solution relying on stochastic signal processing to reduce this effect of noise, producing an estimate of motor adaptation with reduced bias. The result is an improved estimate of trial-by-trial adaptation in a human learner compared to conventional methods. We demonstrate the effectiveness of the new method in analyzing simulated and empirical movement data under different noise conditions. The analytic approach is applicable across different types of movements in varied contexts and should replace the regression analysis method in future motor analysis studies.

**Author Summary:** When a person makes a movement, a motor error is typically observed that then drives motor planning corrections on subsequent movements. This error correction provides insight into how the nervous system is operating, particularly in regard to how much confidence a person places in different sources of information such as sensory feedback or motor command reproducibility. Traditional analysis of movement has required carefully controlled laboratory conditions, limiting the usefulness of motor analysis in clinical and everyday environments. Here we present a new computational method that can be accurately applied to typical movements. Counterintuitive findings of the established approach are corrected by the proposed method. This method will provide a common framework for researchers to analyze movements while extending dynamic motor adaptation analysis capabilities to clinical and non-laboratory settings.

## Introduction

Reaching for a coffee mug, shooting a basketball, or typing on a keyboard - our daily movements are constantly being adjusted based on perceived motor performance. If one misses a basketball shot wide right, the next shot will, on average, end up farther left (hopefully closer to the basket). Adaptation of the motor output of the nervous system is shaped by perceived errors. Measuring the adaptation relationship between changes in motor output and perceived errors provides insight into how humans learn to move and how they rely on different information sources in crafting those movements (1,2).

This adaptation relationship can be calculated by measuring changes in motor output in response to perturbations. However, the corresponding experiments require people to complete movements under abnormal conditions, such as moving through force fields (3–5), moving with shifted visual feedback (6,7), or moving with occasional and unpredictable externally imposed disruptions (8,9). This work has provided significant insight into motor adaptation processes but presents limitations. The experimental manipulations are difficult to implement outside of the laboratory and it remains unclear whether the observed phenomena are relevant to everyday movements. Further, some of these studies require extremely high numbers of repetitive movements.

Trial-by-trial adaptation analysis is an alternative approach to measure this relationship that focuses on error adaptation in unmanipulated environments. This method observes the magnitude of error correction during sequential movements (10–14), and provides similar insight (10) as autocorrelation analysis (15). Trial-by-trial analysis can be run on any type of goal-directed movement under real-world conditions, not relying on applied perturbations. The resulting trial-by-trial adaptation rate has been thought to provide an intuitive metric capturing important motor system characteristics, specifically the relative trust level in control signal generation versus sensory observation (16). Furthermore, empirical observations in this paradigm have been extensively described with computational models including state-space models (17,18) and Bayesian models (1,11,14).

Calculation of trial-by-trial adaptation, however, is easily corrupted by unmeasurable noise within the control signal (19,20), leading to estimates of adaptation that are qualitatively counterintuitive and quantitatively biased. For example, if a person knows their motor output has more noise, they should adapt less. But estimations of trial-by-trial adaptation produce calculations suggesting that they adapt more (11). To date, approaches to estimate trial-by-trial adaptation rates have used an autocorrelation analysis (21,22) or a linear regression analysis (10,11,14) that ignored, by averaging, the effect of stochastic variables at play in a motor system, namely the motor control noise and sensory feedback noise (23). As we describe below, these techniques are sensitive to the noise in the system, such that they produce counterintuitive and biased estimations. The theoretical promise of calculating trial-by-trial adaptation remains, and in this work we propose a novel method to reduce the inherent bias in estimates of the trial-by-trial adaptation rate.

Since it is impossible to externally measure motor noise on a trial-by-trial basis, we sought to estimate the true adaptation rate indirectly. In this work we first demonstrate that the conventional calculation of trial-by-trial adaptation rate, even under steady-state conditions, produces paradoxical results likely due to the overlying noise. Then we present a novel approach to estimate an unbiased trial-by-trial adaptation rate using a model-free analytic stochastic method to filter out overlapping noise. We compare the conventional and proposed method using simulated and empirical data collected with a simple movement experiment. The proposed analytic technique provides a closer approximation to the elusive motor parameters of interest, providing a more useful error adaptation measurement that is relevant for unperturbed movements across a variety of contexts. We conclude with suggestions for how to use the new analysis tool and discuss the wide-ranging research and clinical applications in supporting informative motor assessment.

## Results

Trial-by-trial Adaptation Rate (AR) is broadly defined as the ratio between the trial-to-trial change in movement endpoint and trial error:

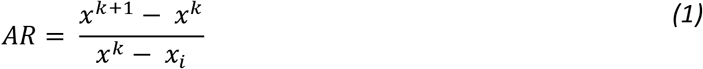

where superscript *k* denotes a given trial number and *x_i_* denotes the intended target. Not that the target modality may be any continuous variable such as position, force, sound frequency, etc. We use the term position throughout the manuscript, but the concepts apply to any signal modality. The generic position (*x*) terms do not adequately differentiate between the various positions, which include the intended position (*x_i_*), the motor command result in a noise-free system (*x_u_*), the externally measured position (*x_m_*), the sensed position (*x_s_*), and the perceived position (*x_p_*) (Fig. 1). See Table I for an overview of variable definitions.

**Fig 1.**
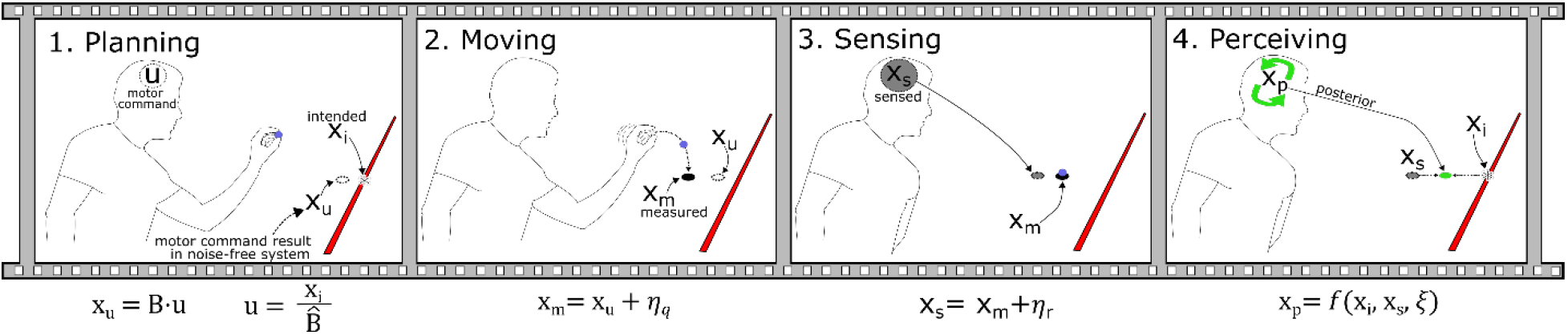
Overview of movement generation framework during a task of tossing a ball to hit a target. In the Planning phase, the thrower generates a motor command (*u*) that, in a noise-free environment, will result in specific ball landing point (*x_u_*). In other words, a control action *u* is formed using an inverse model of the user’s estimate of system dynamics 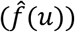, and *x_u_* is obtained by propagating this action *u* through the actual dynamics *f*: 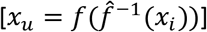. The difference between *x_u_* and the intended target (*x_i_*) represents misestimation of system parameters that are continually updated through the learning process. In the Movement phase, the throw is completed with *x_u_* being affected by *control noise* (*η_q_*) to produce the actual measurable error (*x_m_*). In the Sensing phase, the actual movement endpoint (*x_m_*) is corrupted by feedback noise (*η_r_*), resulting in the endpoint sensed by the thrower (*x_s_*). In the Perceiving phase, a posterior estimate (*x_p_*) of the landing point is arrived at by combining information from the intended endpoint (*x_i_*), the sensed endpoint (*x_s_*), and the level of internal model noise (*ξ*) (1,11).

**Table I.**
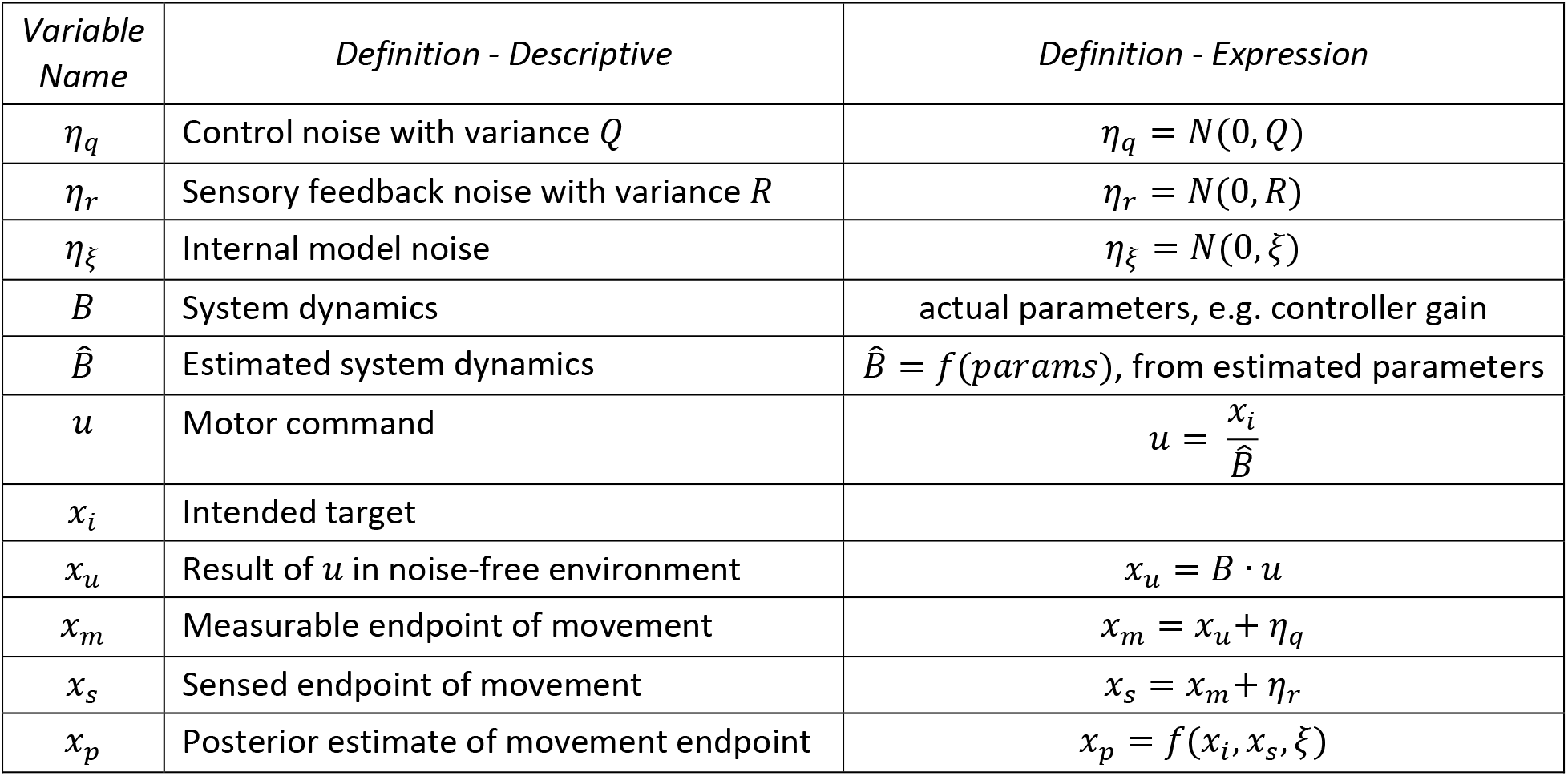
Summary of variable definitions.

When the various types of position are delineated, we can better articulate the implicit assumption that adaptation refers to the response of the person’s intent or noise-free actions (e.g. *x_u_* domain), in response to their perceived error (e.g. *x_p_* domain). We may accordingly use these domains within our adaptation equation. Although impossible to measure empirically, this definition serves as our *gold* standard, defined as:

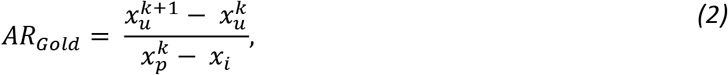

Pragmatically, neither the *x_u_* domain nor the *x_p_* domain can be measured, and so conventional definitions use the *x_m_* domain as a proxy for both:

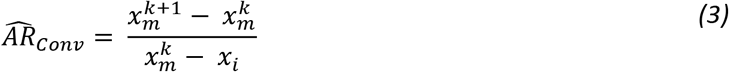

This rate is typically calculated using a linear regression (10,14,24) or auto-correlation (21,22).

To compare the gold standard (2) with the conventional regression analysis (3), we generated simulated movement data across a range of simulation parameters using a Bayesian model of an experienced performer in which trial-by-trial adaptation depends on the subject’s true knowledge of control noise, sensory noise, and internal model confidence (see Methods) (Fig. 2). For increasing control noise (Fig. 2a), the modelled learner should adapt less. The gold-standard definition accurately captures this phenomenon (Fig. 2a, black line). Surprisingly, the conventional estimation shows a qualitatively different phenomenon – its estimate increases as control noise increases (Fig. 2a, red line). For increasing sensory noise (Fig. 2b), the qualitative trend between the two metrics is the same, though the quantitative bias is substantial and qualitatively wrong (an adaptation rate greater than 1 indicates overcompensation).

**Fig 2.**
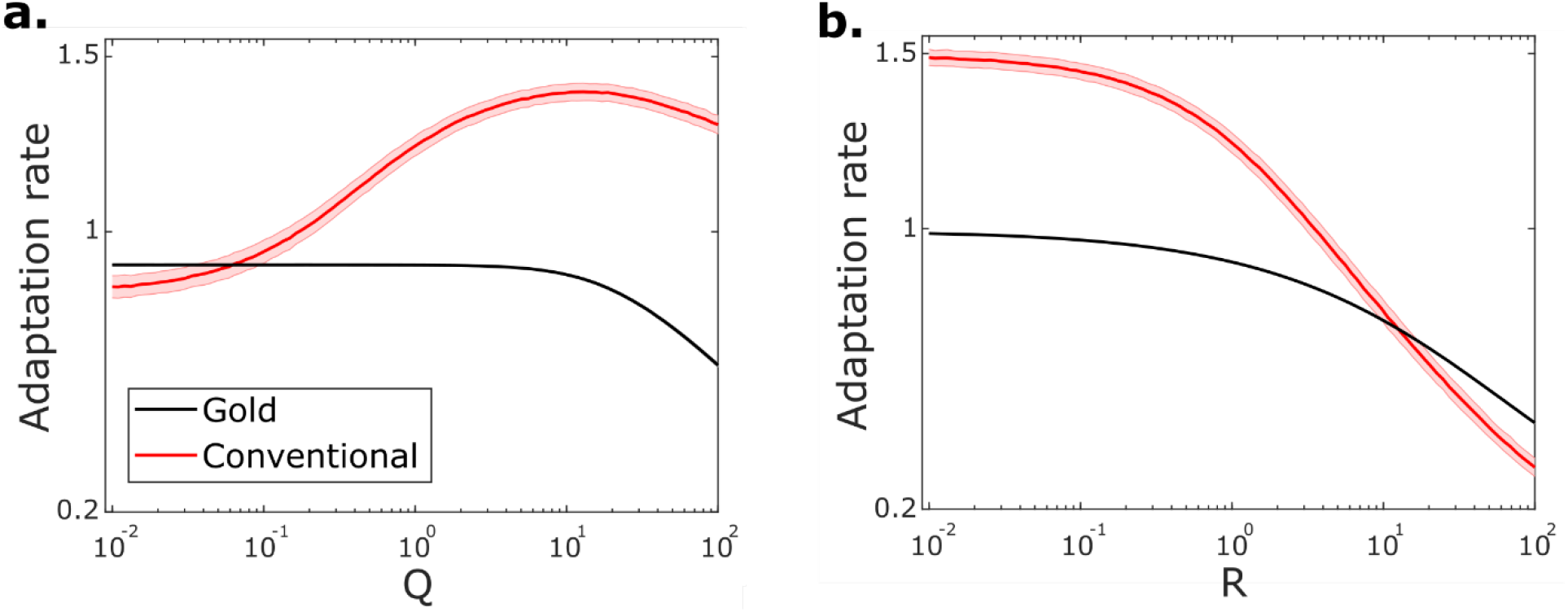
Conventional trial-by-trial adaptation does not capture the expected dynamics of human motor performance. (*a*) Simulated adaptation rate values with changing control noise (Q). The shaded area indicates one standard deviation above and below the mean (solid line). Results from 1,000 simulations at each of 100 values of Q across the range indicated with *x_i_* = 100, *R* = 1 and *ξ* = 0.01. (*b*) Simulated adaptation rate values with changing sensory noise (R). Simulation settings and parameters as in *a* except here *Q* = 1. All adaptation rates are shown as absolute values.

The gold standard definition and conventional estimation differ in the domains used in both their numerator and denominator, and this counterintuitive trend could be caused by either one or a combination of both. As we analytically explain below, using the proxy for the numerator does result in qualitative and quantitative problems. In contrast, using the proxy for the denominator does not produce counterintuitive effects (increases in *η_q_* or *η_r_* lead to increases in estimated adaptation rate, and increases in lead to decreases in estimated adaptation rate, as we would expect) and only mild biases. It is thus important not to use the proxy in the numerator, but less important to avoid the proxy in the denominator. Pragmatically, there is no way to recover *x_p_* from *x_m_* without relying on modelbased assumptions, but we can recover *x_u_* from *x_m_* using simple stochastic analysis, as we show below.

We accordingly define a silver-standard proxy for the adaptation rate as follows:

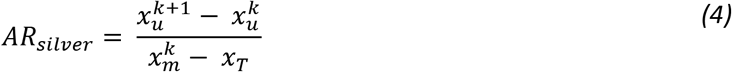

This *silver* definition closely tracks the gold adaptation rate when applied to simulated data with increasing control noise (Fig. 3a) and sensory noise (Fig. 3b). This definition can be calculated under carefully controlled experimental conditions where the control noise added on each trial is known in order to calculate *x_u_* = *x_m_* – *η_q_*, which is the case in the empirical motor study included in this work. The silver adaptation rate can accordingly be used as a baseline against which to compare conventional estimation along with the proposed new estimation approach.

**Fig 3.**
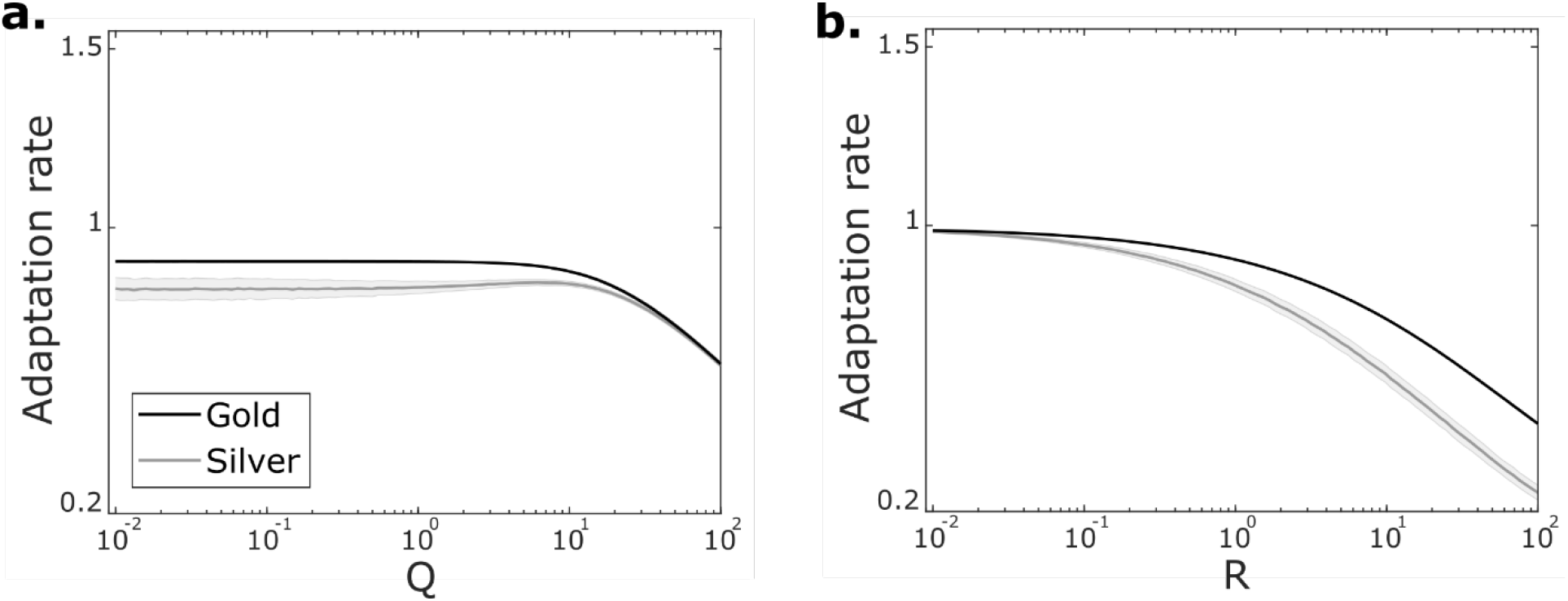
Silver adaptation rate approximates gold standard adaptation rate. Simulated data for changing control noise (*a*) and sensory noise (*b*). Silver can be calculated from data collected in carefully controlled experiments (4). Simulation parameters as in Fig. 2.

The silver definition provides an experimentally measurable baseline that is in qualitative and quantitative agreement with the gold standard, but it cannot be measured in real-world conditions without applying perturbations – the very conditions that make trial-by-trial adaptation so appealing. We accordingly sought to develop a way to estimate the true adaptation rate that could be applied to a variety of laboratory and real-world movement data. The analytic approach, using statistical principles to filter out the noise effects and estimate the silver adaptation rate, can be defined as follows (see Methods for derivation):

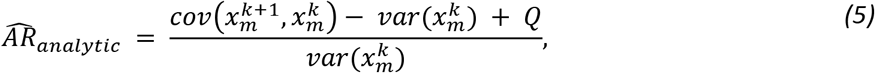

where the variance of control noise *Q* can be measured or estimated from the experiment. We systematically varied simulation parameters (as in Fig. 2) and compared the analytic estimate to the gold AR, silver AR and the conventional regression estimate (Fig. 4).

**Fig 4.**
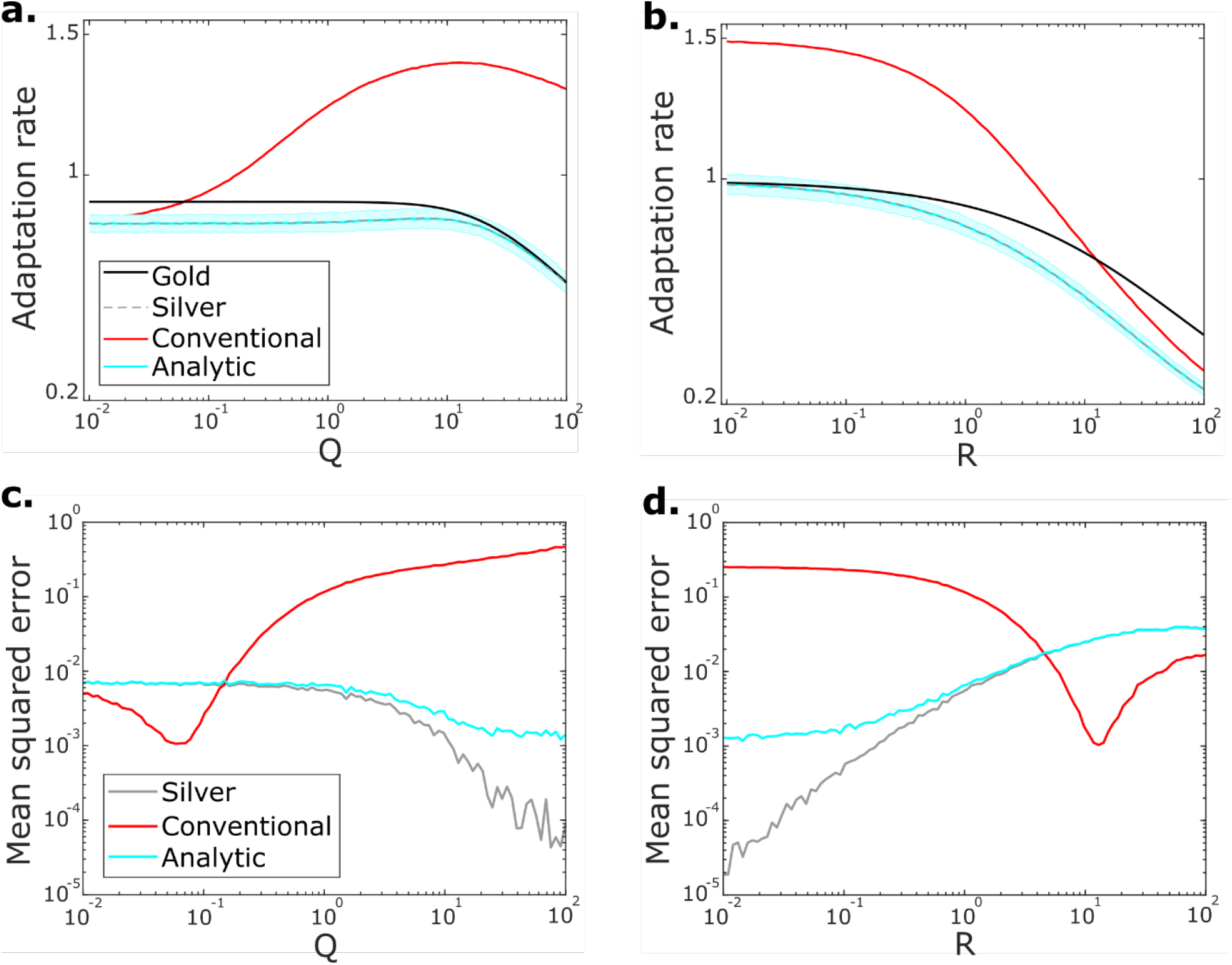
Comparison of different trial-by-trial adaptation rate calculation methods. (a-b) Simulation and parameters settings as in Fig. 2 for changing control noise with R = 1 (a) and changing sensory noise with Q = 1 (b). For clarity, the one standard deviation shaded range is shown only for the analytic results (see Fig. 2 and 3 for variability shading for other analysis methods). (c-d) Performance of each analysis method shown in (a) and (b) as measured using the mean squared error compared to the gold adaptation rate for changing control noise (c) and sensory noise (d).

The analytic estimate better captures the qualitative and quantitative trends observed for changing control noise compared to the conventional regression estimate (Fig. 4a). We also observed more quantitatively aligned estimates under conditions of changing sensory noise (Fig. 4b). Estimation error plotted across the changing parameters shows that the analytic estimate consistently produces better estimates of the gold adaptation rate than the conventional regression method (Fig. 4cd). The one exception is that due to the different trends in adaptation rates with changing parameters, the conventional regression happens to overlap with the gold adaptation rate showing a low estimation error indicated by low mean squared error when Q<R (Fig. 4cd). When internal model noise (*ξ*) was varied we observed similar quantitative improvements using the new method compared to the conventional regression estimate.

Estimation results presented so far involved the analysis of 1,000 trials of simulated movement data, a number not always feasible for everyday experiments. We sought to determine the variability of the estimation methods with different numbers of trials analyzed. We systematically varied the size of the analyzed trial window and observed the performance of each analysis technique on simulated data (Fig. 5). As the number of trials analyzed increases, the variability of each estimation method is reduced (Fig. 5).

**Fig 5.**
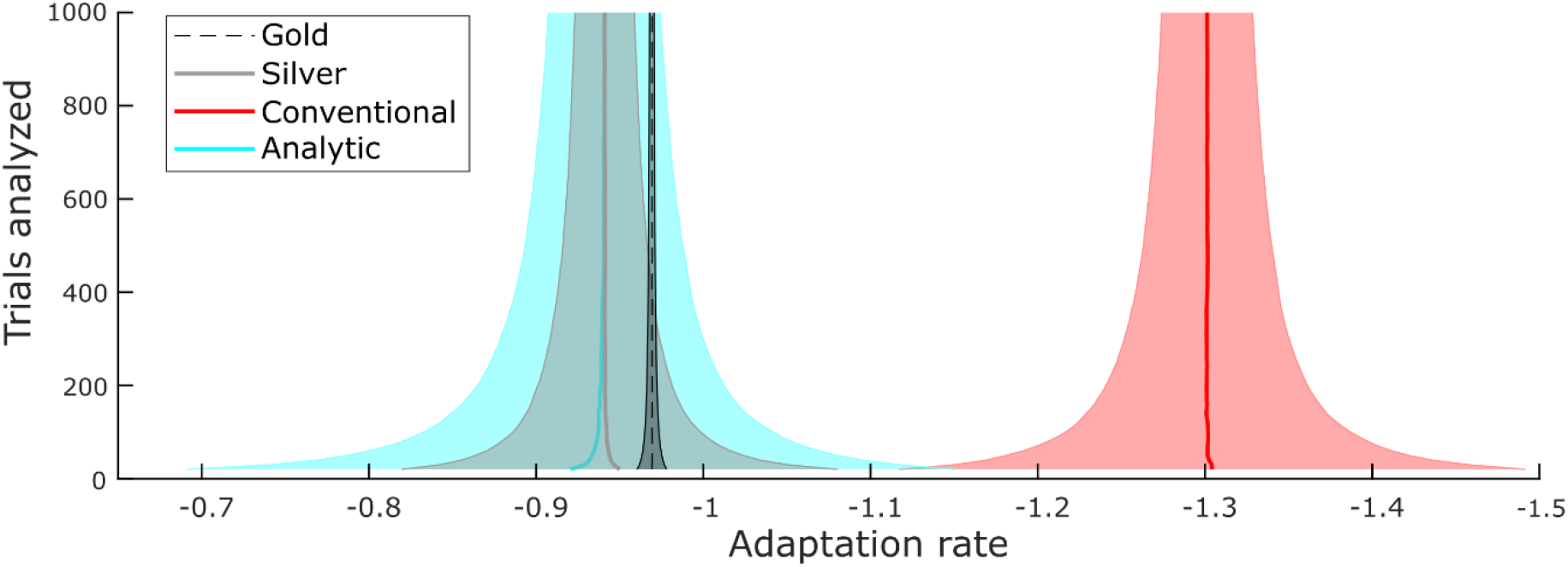
Analyzing more trials reduces variability of results. 10,000 simulations were run with the following parameters: *Q* = 1, *R* = 1, *ξ* = 0.1. Different windows of trials were analyzed on each simulation run with the shaded area indicating one standard deviation of the total results.

We observed a similar improvement in estimated adaptation rates using the analytic method in an empirical study (Fig. 6). Twenty-seven able-bodied participants each completed three sets of 100 movements under different noise conditions. Participants moved a mouse cursor on a screen to a target and were provided with endpoint landing position feedback only. The order of the three noise conditions – NO added noise, LOW added noise, HIGH added noise – was randomized for each participant. The endpoint error and externally applied noise were recorded for every trial and then used to run the adaptation rate estimation methods.

**Fig 6.**
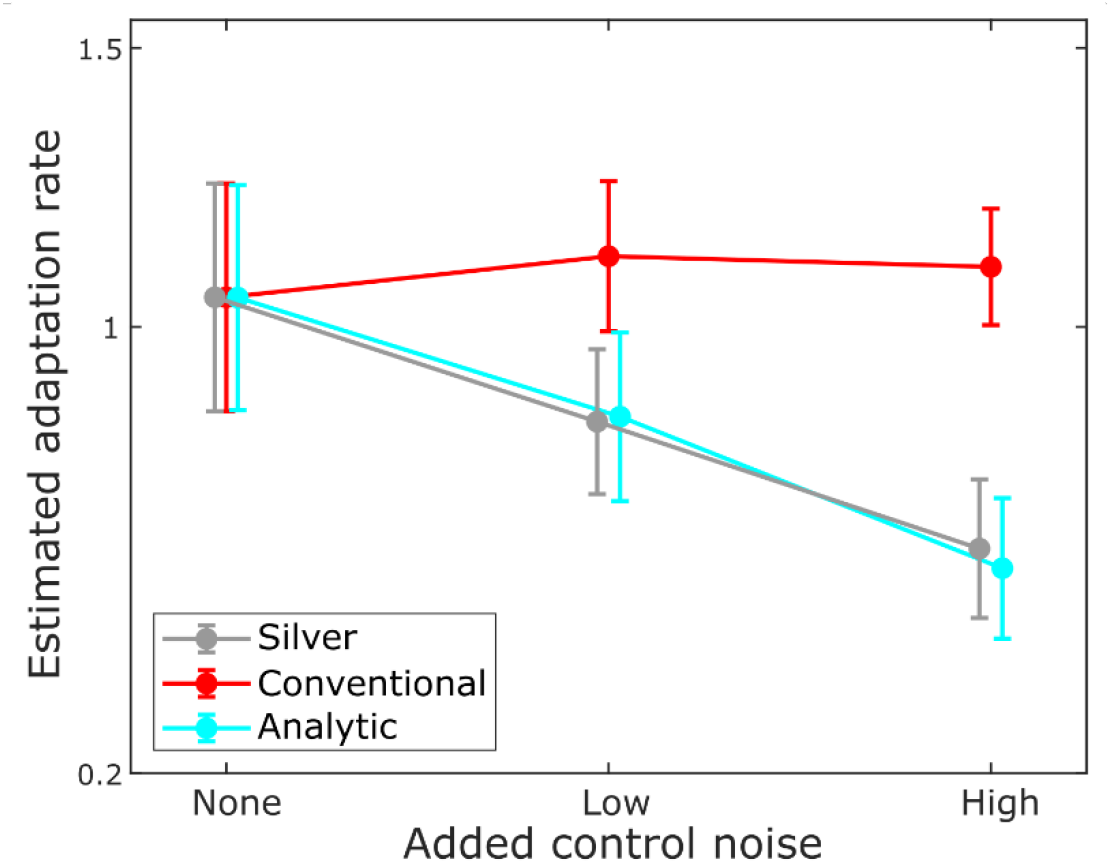
Performance of analysis techniques on empirical data. The means with standard deviation error bars comparing the three analysis methods run on data collected from 27 able-bodied participants completing a computer cursor movement study under different added noise conditions. Note that the silver adaptation rate and conventional estimate are mathematically equivalent with no added control noise. All pairwise differences within silver ARs and analytic ARs are statistically significant (ANOVA with Bonferroni-corrected post-hoc comparison, p<.001).

Increasing levels of control noise resulted in a slightly increasing trend in conventional regression estimates of trial-by-trial adaptation using the conventional regression approach (Fig. 6, red line), matching the observations on the simulated datasets (Fig. 2a). However, as predicted, the silver adaptation rate estimate showed a proportional decrease with increasing control noise (Fig. 6, gray line), a trend closely matched by the analytic estimate (Fig. 6, cyan line).

The data resulting from each analysis method shown in Fig. 6 were analyzed with a separate one-way ANOVA with repeated measures using Bonferroni corrected post-hoc comparisons:

- For the conventional estimates of trial-by-trial adaptation rate, Mauchly’s Test of Sphericity indicated that the sphericity assumption was not violated (χ^2^(2) = 2.595, p = .273). The repeated measures ANOVA did not indicate statistical significance between estimated adaptation rates across different noise conditions (F(2,52) = 2.389, p = .102).
- For the silver trial-by-trial adaptation rates, Mauchly’s Test of Sphericity indicated that the sphericity assumption was violated (χ^2^ (2) = 8.208, p = .017). The repeated measures ANOVA with Greenhouse-Geisser correction indicated a statistically significant difference of estimated adaptation rates across different noise conditions (F(1.563,40.629) = 68.480, p < .001). All Bonferroni-corrected pairwise comparisons were statistically significant (p < .001).
- For the analytic estimates trial-by-trial adaptation rate, Mauchly’s Test of Sphericity indicated that the sphericity assumption was not violated (χ^2^(2) = 2.777, p = .249). The repeated measures ANOVA indicated a statistically significant difference of estimated adaptation rates across different noise conditions (F(2,52) = 73.551, p < .001). All Bonferroni-corrected pairwise comparisons were statistically significant (p < .001).

In summary, the conventional adaptation method was not able to capture the clear decrease in adaptation caused by increasing control noise (silver), whereas the analytic method was able to do so, and its estimate closely aligned with the silver definition.

## Methods

In the following subsections we first present the derivation of the analytic estimation approach and then describe the simulations and experiments we used to assess our techniques.

### Analytic estimation of adaptation rate

The least squares regression of the slope *β* for the linear relationship *y* = *α* + *βx* is:

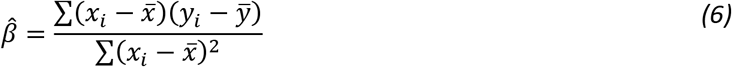

However, the contribution of noise sources in *x* and *y* are not apparent from this expression, and as we have shown in the results, it is important to be able to compensate for noise sources in *y*. An equivalent well-known analytic representation may be used, which makes use of stochastic methods to produce a probabilistic estimator:

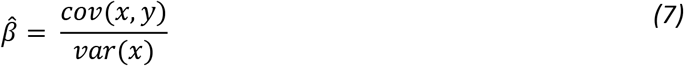

It will be easier to see and remove the contribution of noise sources using this analytic estimation. Applying this analytic estimate of the slope to our silver definition of adaptation rate (4), our estimate of AR is:

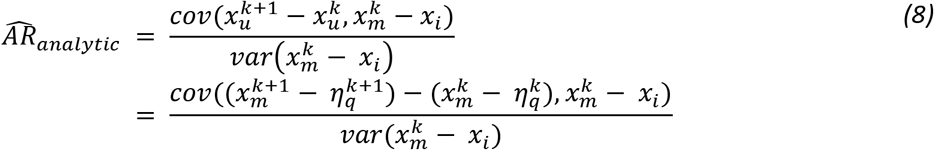

Simplifying the numerator,

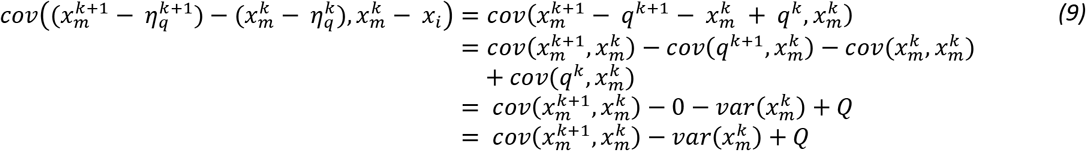

Simplifying the denominator,

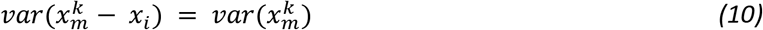

Combining our simplified numerator and denominator, our estimate accordingly can be calculated as:

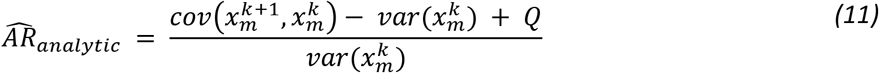

We call Eq. (11) the *analytic* estimate of the trial-by-trial adaptation rate. It can be calculated from endpoint error records of movements and an estimate of the control noise variance.

### Simulations

#### Simulated data generation

For all simulated data, MATLAB software (MathWorks, Natick, MA, version 2020a) was used to run a hierarchical Kalman filter model that has been described in detail elsewhere (10). Briefly, the first part of the model uses a Kalman filter to generate a posterior estimate of the endpoint position (i.e. *x_p_*) by fusing the a priori premotor estimate (i.e. *x_i_*) and the post-movement sensory observation (i.e. *x_s_*). The second layer of the model uses another Kalman filter to generate an update in the learner’s estimation of system parameters. In this case we use a single parameter representing the gain of the controller; the learner’s misestimation of the controller gain leads to motor errors. The magnitude of the parameter estimate update is determined by the second Kalman filter’s integration of the perceived error and the overall internal model uncertainty that is driven by the uncertainty increment at each trial (*ξ*). For example, if the internal model uncertainty is high, parameter estimate updates will be larger.

Simulated data were generated with changing input parameters to test the sensitivity of each estimation method. Each of the three input parameters – *Q, R*, and *ξ* – was systematically varied while the other two were held constant. Constant parameters values were as follows: *Q* = 1, *R* = 1, = 0.1, *x_i_* = 100. When varied, 100 equally spaced values on a log scale were used for the changing input parameter ranging from 10^-2^ to 10^2^, enabling the parameter to range from being dominated to dominating. For each set of input parameters, 1,000 simulations were run, each with 1,500 movement trials with the last 1000 trials used for analysis. All other parameters were constant including the controller gain (*B* = 1), initial estimated gain 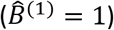, and the initial overall internal model uncertainty 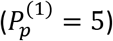. The gold, silver, conventional, and analytic adaptation rate estimates x1were calculated for each simulated dataset. Mean squared errors were computed comparing the estimates to the gold trial-by-trial adaptation rate.

To explore the sensitivity of the estimation methods when run on different numbers of trials, a separate set of simulated data was generated with fixed parameters (*Q* = 1, *R* = 1, *ξ* = 0.1). For each of 10,000 simulation runs, data from 2000 movement trials were generated. Starting at the 1000^th^ trial, the data were analyzed using each estimation procedure across 99 trial windows of varying sizes, equally spaced from 20 trials to 1000 trials.

### Empirical study

#### Ethics Oversight

All research with human participants was conducted with approval and oversight by the Rhodes College Institutional Review Board. All participants provided written informed consent.

#### Participants

27 able-bodied and right-handed participants [mean age = 18.7yrs, range = 18-21, 23 females], recruited from the Rhodes College Psychology Department participant pool, completed the experiment. Two additional participants did not complete the full experiment and their data were not included. Participants were compensated for their time with class credit points.

#### Setup

While seated comfortably at a desk and viewing a 27” computer monitor (Acer Model #G276HL), participants were asked to move a computer cursor from left to right along a straight line to hit a target at a distance of 22.4cm. Participants used a wireless mouse (Logitech Model #G703) with a reduced sensitivity setting on an extra wide mouse pad (31.5” x 11.8”) to land on a stationary onscreen target. The movement was initiated with a participant mouse click that caused the cursor to disappear until the participant completed the movement and clicked again. At the completion of each movement, endpoint only visual feedback was provided on screen. View of the participant’s hand and arm movement was blocked by a rigid covering that did not contact the participant or impede movement of the computer mouse.

Participants wore noise canceling headphones with white noise playing to mask ambient auditory information. Participants were alerted with on-screen text when movement endpoints were off-screen, backwards or too short (landing <25% of the way to the target).

All participants completed three blocks of 100 successful trials, each under different noise conditions. Each movement block took about 6 minutes to complete. In one of the three blocks, movements were completed without added noise. In the Low noise block, control noise of variance 5.04 cm^2^ was added, with noise of 20.11 cm^2^ variance added in the High noise block. The noise was applied by computing a random number from a Gaussian distribution with the variance for that movement block and adding that shift to the landing position before the cursor was displayed to the participant. Each participant was randomly assigned one of six possible orders for the three noise conditions to appear.

To encourage participant engagement, a scoring system was implemented. Scores were displayed on-screen with points awarded for movements close to the target. 200 points were awarded for direct target hits, 100 points for movement endpoints within 25 pixels (0.78cm) of the target, and 50 points for within 100 pixels (3.12cm) of the target.

#### Analysis

To prepare the data for analysis, backwards and short movements were presumed to be accidental mouse clicks and removed from the movement records. Error during off-screen movements could not be measured and this resulted in a break in contiguous analyzable movement trials. The longest stretch of contiguous movement trials without an off-screen movement was extracted for each block of data to be analyzed. Steady-state trials in which initial parameter learning had stabilized were identified from the contiguous movement trials using a previously reported method (10). Across the 81 blocks of data collected, the average number of steady state trials analyzed was 93. Only 3 movement blocks resulted in fewer than 67 steady state trials. For one block which resulted in only 11 steady state trials identified, the last 20 trials were analyzed. As in the simulated data, contiguous movement trials were analyzed using the conventional (3), silver (4), and analytic (5) methods. For the silver method, the added noise on each trial was used, ignoring any baseline control noise generated by the participant. Likewise, for the analytic method the added control noise variance (Q) was used.

### Data and code availability

Code used to simulate data, run the estimation analyses, and generate the figures, along with the simulated data used for estimation method analysis, and the empirical data are available here: https://osf.io/4vsmd/?view_only=9fe78f28eefe4a08aafa56e84cbd9397 (permanent DOI pending publication).

## Discussion

Here we have developed an analytic method to better estimate trial-by-trial adaptation rates. When applied to simulated data, the novel approach produces qualitatively accurate and quantitively improved estimates of the gold adaptation rate, compared to the conventional regression estimate. Following validation with simulated data, we ran an empirical study in which participants moved a cursor on a screen with a computer mouse under varied noise conditions. Analysis of empirical data matched the simulation results: the novel analysis approach produced better adaptation rate estimates compared to the conventional regression estimate. In the empirical work, adaptation estimates were compared to a close approximation of the gold adaptation rate – the silver estimate – a value that can only be calculated under carefully controlled experimental conditions. The analytic adaptation rate estimation method proposed here can be run across a wide range of motor contexts, both within and outside of the laboratory. Due its versatility, accuracy and sensitivity, the new proposed technique provides a path to reduce bias in the analysis of human motor performance.

The estimation method resulting from this work can be applied in the laboratory to advance motor adaptation research in several ways. First, the method provides a well-justified and standardized approach to measure adaptation. In the past there have been different ways to measure adaptation that result in incomparable numerical indicators, e.g. trial-by-trial adaptation and perturbation adaptation (10). The new method can be applied in a wide range of contexts to allow for direct comparisons across different experimental setups. Second, the method reduces experimental limitations associated with perturbation adaptation studies. These perturbation studies, looking at the magnitude of adaptation to an unexpected externally applied disturbance, avoid some noise biasing associated with trial-by-trial adaptation because the perturbation magnitude dominates the baseline system noise. One issue is that the applied perturbations can alter motor dynamics, interfering with the process being observed. Perturbation adaptation studies also require large numbers of trials because the disturbances can only occur sporadically. The new method eliminates both issues associated with perturbation adaptation studies.

The application of this novel approach to estimate adaptation rate goes beyond the laboratory environment. Since the method operates on unperturbed movement data, it can be run in clinical or real-world settings. For example, a stroke patient could make repetitive reaching movements with an occupational therapist. Understanding the magnitude of error adaptation is critically insightful in such a case, providing details about motor deficiencies and compensatory strategies that are overlooked by currently available clinical motor assessments. Or movement data could be extracted from video recordings of a pitcher in a baseball game; without the need for manipulations or complicated measurements, the adaptation analysis approach can be applied to almost any sequence of movements.

Although the analytic method provides accurate estimates, it does require knowledge of the baseline control noise (Q). This parameter can be measured with a separate experiment in which the learner makes movements in the absence of sensory feedback and the endpoint variability is recorded. Sporadic feedback is necessary to reduce endpoint drift and although removing exteroceptive feedback is straightforward, proprioceptive feedback usually remains. There are other ways to estimate control noise including measurement of the just-noticeable-difference of a perturbation that can be converted to a total noise estimate (25) with subsequent subtraction of an estimate of sensory noise. Another limitation of the analytic method is that its results show high variability similar to the conventional regression method when low numbers of trials are analyzed (Fig. 5). The improved estimation accuracy is maintained across trial window sizes and we suggest analyzing as many trials as possible.

A few limitations of the validation of the proposed protocol should be addressed. The need for simulated data to test and compare the estimation methods requires the adoption of a specific motor control model that requires assumptions about the underlying computational mechanisms driving motor processes. Here we used a Bayesian model of motor control which has been shown to be supported in many (1,14) but not all motor contexts (16). Other models could have been used including state-space only models (17,18) or cost-function models (26–28). Nevertheless, the estimation improvements we observed in the empirical data were similar to those observed with simulated data, suggesting the assumptions required to generate simulated motor data have not affected our findings. The primary motivation was in overcoming *qualitative* issues with the conventional regression estimation, further emphasizing that the specific model chosen is not critical. Even if the underlying motor control model to generate simulated data were different, the estimation methods should produce qualitatively similar results. Any effect on the results associated with choosing a different model would be represented by a consistent shift in quantitative results, something we may expect when going from simulated to empirical data anyway.

In collecting empirical data to test the proposed method, a few limitations arose. Since we wanted a precise estimate of control noise and we were comparing against the silver estimate that requires knowledge of the noise added to each movement trial, we focused only on the noise we experimentally added. This approach ignored the baseline control noise inherent in any moving human. Since we were most interested in qualitative trends, we were justified in ignoring baseline noise for the silver and analytic approaches. It is not possible to ignore the baseline noise in the conventional regression approach but since the noise we added was much greater than baseline noise, we were still able to observe and compare qualitative trends.

Another limitation was the possible impact of order effects in our within-subjects experimental design. We randomized the order of conditions for each participant to average out any order effect, with the consequence of potentially noisier results. And finally, differences in sensory noise between participants could have affected the results. We tried to eliminate as much feedback as possible by using noise reducing headphones and by visually occluding the moving arm, but proprioceptive inputs were not eliminated. The ability to interpret proprioceptive inputs to drive motor adaptation will vary for each individual research participant, leading to noisier empirical data that may be differentially impactful across control noise conditions. Again, since the focus was on qualitative trends, these individual differences should not affect the conclusions of the study. In our simulations we were able to keep sensory noise constant and we observed similar results. If sensory noise is a concern to others using these methods, it can be measured using a separate experimental protocol (29).

Here we have demonstrated that current motor adaptation analysis is biased by noise, and we provide an important methodological advance to correct this issue. The result is a more accurate estimation of trial-by-trial adaptation that better captures what makes this metric useful: the adaptation of *motor intent* based on perceived error. Not only will the analytic adaptation rate estimation solution we provide support improved laboratory analysis and the correction of possible misinterpretation of data in previous studies, but it will allow for the analysis of a wide range of motor behavior in real-world settings. Using trial-by-trial adaptation rates as an informative clinical motor metric is promising, and now possible with the advances presented here.

## Acknowledgements

We gratefully acknowledge helpful conversations and suggestions by Phil Parker on the topic of stochastic estimation. The work by AS was funded by the Sensory Motor Adaptive Rehabilitation Technology (SMART) Network and the Smart Technology (ST) Innovations at the University of Alberta.

